# Task-dependent and automatic tracking of hierarchical linguistic structure

**DOI:** 10.1101/2022.02.08.479571

**Authors:** Sanne Ten Oever, Sara Carta, Greta Kaufeld, Andrea E. Martin

## Abstract

Linguistic phrases are tracked in sentences even though there is no clear acoustic phrasal marker in the physical signal. This phenomenon suggests an automatic tracking of abstract linguistic structure that is endogenously generated by the brain. However, all studies investigating linguistic tracking compare conditions where either relevant information at linguistic timescales is available, or where this information is absent altogether (e.g., sentences versus word lists during passive listening). It is therefore unclear whether tracking at these phrasal timescales is related to the content of language, or rather, is a consequence of attending to the timescales that happen to match behaviourally-relevant information. To investigate this question, we presented participants with sentences and word lists while recording their brain activity with MEG. Participants performed passive, syllable, word, and word-combination tasks corresponding to attending to rates they would naturally attend to, syllable-rates, word-rates, and phrasal-rates, respectively. We replicated overall findings of stronger phrasal-rate tracking measured with mutual information (MI) for sentences compared to word lists across the classical language network. However, in the inferior frontal gyrus (IFG) we found a task-effect suggesting stronger phrasal-rate tracking during the word-combination task independent of the presence of linguistic structure, as well as stronger delta-band connectivity during this task. These results suggest that extracting linguistic information at phrasal-rates occurs automatically with or without the presence of an additional task, but also that that IFG might be important for temporal integration across various perceptual domains.

## Introduction

Understanding spoken language requires a multitude of processes [1–3]. Acoustic patterns have to be segmented and mapped onto internally stored phonetic and syllabic representations [3–5]. These phonemes have to be combined and mapped onto words which then have to be mapped to abstract linguistic phrasal structures [2, 6]. Proficient speakers of a language seem to do this so naturally that one might almost forget the complex parallel and hierarchical processing which occurs during natural speech and language comprehension.

It has been shown that it is essential to track the temporal dynamics of the speech signal in order to understand its meaning [7, 8]. In natural speech, syllables follow up on each other in the theta range (3-8 Hz; [9–11]), while higher-level linguistic features such as words and phrases occur at lower rates (0.5-3 Hz; [9, 12, 13]). Tracking of syllabic features is stronger when one understands a language [14–16] and tracking of phrasal rates is more prominent when the signal contains phrasal information ([12, 13, 17]; e.g., word lists versus sentences). Importantly, phrasal tracking even occurs when there are no distinct acoustic modulations at the phrasal rate [12, 13, 17]. These results seem to suggest that tracking of relevant temporal timescales is critical for speech understanding.

An observation one could make regarding these findings is that tracking occurs only at the rates that are meaningful and thereby behaviourally relevant [12, 17]. For example, in word lists, word-rate is the slowest rate that is meaningful during natural listening. Modulations at slower phrasal rates might not be tracked as they do not contain behaviourally relevant information. In contrast, in sentences phrasal rates contain linguistic information and therefore these slower rates are also tracked. Thus, when listening to speech one automatically tries to extract the meaning, which requires extracting information at the highest linguistic level [3, 5]. However, it is unsure if the tracking at these slower rates is a unique feature of language processing or rather dependent on the level of attention to relevant temporal timescales.

As understanding language requires a multitude of processing, it is difficult to figure out what participants actually are doing when listening to natural speech. Moreover, designing a task in an experimental setting that does justice to this multitude of processing is difficult. This is probably why tasks in language studies vary vastly. Tasks include passively listening (e.g. [12], asking comprehension questions (e.g. [13], rating intelligibility (e.g. [14, 16], working memory tasks (e.g. [18], or even syllable counting (e.g. [17]. It is unclear whether outcomes are dependent on the specifics of the task. There has so far not been a study that investigates if task instructions focusing on extracting information at different temporal rates or timescales have an influence on the tracking that occurs on these timescales. It is therefore not clear whether tracking phrasal timescales is unique for language stimuli which contain phrasal structures, or could also occur for other acoustic materials where participants are instructed to pay attention to information happening at these temporal rates or timescales.

To answer this question, we designed an experiment in which participants were instructed to pay attention to different temporal modulation rates while listening to the same stimuli. We presented participants with naturally spoken sentences and word lists and asked them to either passively listen, or perform a task on the temporal scales corresponding to syllables, words, or phrases. We recorded MEG while participants performed these tasks and investigated tracking as well as power and connectivity at three nodes that are part of the language network: the superior temporal gyrus (STG), the middle temporal gyrus (MTG), and the inferior frontal gyrus (IFG). We hypothesized that if tracking is purely based on behavioural relevance, it should mostly depend on the task instructions, rather than the nature of the stimuli. In contrast, if there is something automatic and specific about language information, tracking should depend on the level of linguistic information in the acoustic signal.

## Methods

### Participants

In total twenty Dutch native speakers (16 females; age range: 18-59; mean age = 39.5) participated in the study. All were right-handed, reported normal hearing, had normal or corrected-to-normal vision, and did not have any history of dyslexia or other language related disorders. Participants performed a screening for their eligibility in the MEG and MRI and gave written informed consent. The study was approved by the ethical Commission for human research Arnhem/Nijmegen (project number CMO2014/288). Participants were reimbursed for their participation. One participant was excluded from the analysis as they did not finish the full session.

### Materials and design

Materials were identical to the stimuli used in Kaufeld et al., [12]. They consisted of naturally spoken sentences or word lists which consisted of 10 words (see Table 1 for examples). The sentences contained two coordinate clauses with the following structure: [Adj N V N Conj Det Adj N V N]. All words were disyllabic except for the words “de” (*the*) and “en” (*and*). Word lists were word-scrambled versions of the original sentences which always followed the structure [V V Adj Adj Det Conj N N N N] or [N N N N Det Conj V V Adj Adj] to ensure that they were grammatically incorrect. In total sixty sentences were used. All sentences were presented at a comfortable sound level.

**Table 1.**
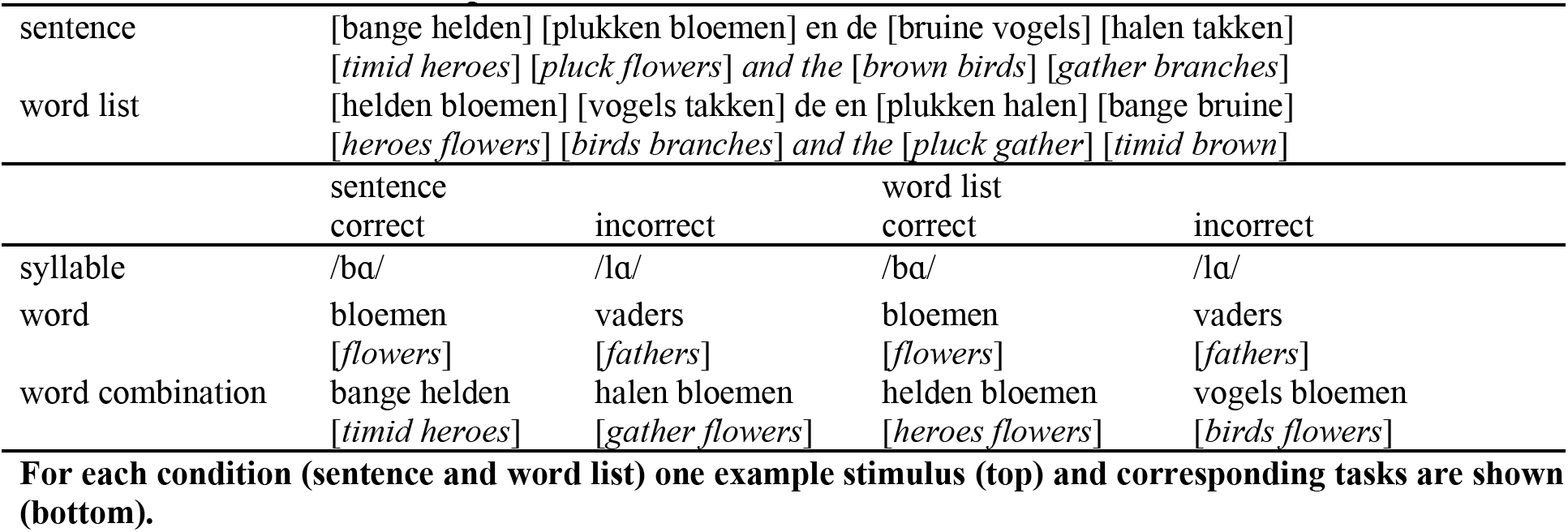
Stimuli and task examples.

Participants were asked to perform four different tasks on these stimuli: a passive task, a syllable task, a word task, and a word combination task. For the passive task, participants did not need to perform any task other than comprehension – they only needed to press a button to go to the next trial. For the syllable task, participants heard after every sentence two part-of-speech sounds, each consisting of one syllable. The sound fragments were a randomly determined syllable from the previously presented sentence and a random syllable from all other sentences. Participants’ task was to indicate via a button press which of the two sound fragments was part of the previous sentence. For the word task, two words were displayed on the screen after each trial (a random word from the just presented sentence and one random word from all other sentences excluding “de” and “en”), and participants needed to indicate which of the two words was part of the sentence before. For the word combination task, participants were presented with two word pairs on the screen. Each of the four words was part of the just presented sentence, but only one of the pairs was in the correct order. Participants needed to indicate which of the two pairs was presented in the sentence before. Presented options for the sentence condition were always a grammatically and semantically plausible combination of words. See Table 1 for an example of the tasks for each condition (sentences and word lists). The three active tasks required participants to focus on the syllabic (syllable task), word (word task), or phrasal (word combination task) timescales.

### Procedure

At the beginning of each trial, participants were instructed to look at a fixation cross presented at the middle of the screen on a grey background. Audio recordings were presented after a random interval between 1.5-3 seconds; 1 second after the end of the audio, the task was presented. For the word and word combination task, this was the presentation of visual stimuli. For the syllable task, this entailed presenting the sound fragments one after each other (with a delay of 0.5 seconds in between). For the passive task, this was the instruction to press a button to continue. In total there were eight blocks (two conditions * four tasks) each lasting about 8 minutes. The order of the blocks was pseudo-randomized by independently randomizing the order of the tasks and the conditions. We then always presented the same task twice in a row to avoid task-switching costs. As a consequence, condition was always alternated (a possible order of blocks would be: passive-sentence, passive-word list, word-sentence, word-word list, syllable-sentence, syllable-word list, word combination-sentence, word combination-word list). After the main experiment, an auditory localizer was collected which consisted of listening to 200ms sinewave and broadband sounds (centred at 0.5, 1, and 2 kHz; for the broadband at a 10% frequency band) at approximately equal loudness. Each sound had a 50ms linear on and off ramp and was presented for 30 times (with random inter-stimulus interval between 1 and 2 seconds).

At arrival, participants filled out a screening. Electrodes to monitor eye movements and heart beat were placed (left mastoid was used as ground electrode) at an impedance below 15 kiloOhm. Participants wore metal free clothes and fitted earmolds on which two of the three head localizers were placed (together with a final head localizer placed at the nasion). They then performed the experiment in the MEG. MEG was recorded using a 75-channel axial gradiometer CTF MEG system at a sampling rate of 1.2 kHz. After every block participants had a break, during which head position was corrected [19]. After the session, the headshape was collected using Polhemus digitizer (using as fiducials the nasion and the entrance of the ear canals as positioned with the earmolds). For each participant, an MRI was collected with a 3 T Siemens Skyra system using the MPRAGE sequence (1mm isotropic). Also for the MRI acquisition participants wore the earmolds with vitamin pills to optimize the alignment.

### Behavioural analysis

We performed a linear mixed model analysis with fixed factors task (syllable, word, and word combination) and condition (sentence and word list) as implemented by lmer in R4.1.0. The dependent variable was accuracy. First, any outliers were removed (values more extreme than median± 2.5 IQR). Then, we investigated what the best random model was, including a random intercept or a random slope for one or two of the factors. The models with varying random factors were compared with each other using an ANOVA. With no significant difference, the model with the lowest number of factors was included (with minimally a random intercept). Finally, lsmeans was used for follow-up tests using the kenward-roger method to calculate the degrees of freedom from the linear mixed model. For significant interactions, we investigated the effect of condition per task. For main effects, we investigated pairwise comparisons. We corrected for multiple comparisons using adjusted Bonferroni corrections. For all further reported statistical analyses for the MEG data, we followed the same procedure (except that there was one more level of task, i.e. the passive task). To avoid exploding the amount of comparisons, we a-priori decided for any task effects in the MEG analysis to only compare the individual tasks with the phrase task.

### MEG pre-processing

First source models from the MRI were made using a surface-based approach in which grid points were defined on the cortical sheet using the automatic segmentation of freesurfer6.0 [20] in combination with pre-processing tools from the HCP workbench1.3.2 [21] to down-sample the mesh to 4k vertices per hemisphere. The MRI was co-registered to the MEG using the previously defined fiducials as well as an automatic alignment of the MRI to the Polhemus headshape using the Fieldtrip20211102 software [22].

Pre-processing involved epoching the data between −3 and +7.9 seconds (+3 relative to the longest sentence of 4.9 sec) around sentence onset. We applied a dftfilter at 50, 100 and 150 Hz to remove line noise, a Butterworth bandpass filter between 0.6 and 100 Hz, and performed baseline correction (−0.2-0 sec baseline). Trials with excessive movements or squid jumps were removed via visual inspection (20.1±18.5 trials removed; mean±standard deviation). Then data was resampled to 300 Hz and we performed ICA decomposition to correct for eye blinks/movement and heart beat artefacts (4.7±0.99 components removed; mean±standard deviation). Trials with remaining artefacts were removed by visual inspection (11.3±12.4 trials removed; mean±standard deviation). Then we applied a lcmv filter to transform the data to have single-trial source space representations. A common filter across all trials was calculated using a fixed orientation and a lambda of 5%. We only extracted time courses for our regions of interest (superior temporal gyrus [1,29,32,33], medial temporal gyrus [6,8,14], and inferior frontal cortex [17,18,19]; numbers correspond to label-coding from the aparc parcellations implemented in Freesurfer). These time courses were baseline corrected (−0.2 to 0 seconds). To reduce computational load and to ensure that we used relevant data within the ROI, we extracted the top 20 PCA components per ROI for all following analyses based on a PCA using the time window of interest (0.5-3.7 seconds; 0.5 to ensure that all initial evoked responses were not included and 3.7 as it corresponds to the shortest trials).

### Mutual information analysis

First, we extracted the speech envelopes by following previous procedures [12, 13, 23]. The acoustic waveforms (third-order Butterworth filter) were filtered in eight frequency bands (100-8000 Hz) equidistant on the cochlear frequency map [24]. The absolute of the Hilbert transform was computed, we low-passed the data at 100 Hz (third order Butterworth) and then down-sampled to 300 Hz (matching the MEG sampling rate). Then, we averaged across all bands.

Mutual information (MI) was calculated between the filtered speech envelopes and the filtered MEG data at three different frequency bands corresponding to information content at different linguistic hierarchical levels: phrase (0.8-1.1 Hz), word (1.9-2.8 Hz), and syllable (3.5-5.0 Hz). Our main analysis focusses on the phrasal band, as that is where our previous study found the strongest effects [12], but for completeness we also report on the other bands. Mutual information was estimated after the evoked response (0.5 sec) until the end of the stimulus at five different delays (60, 80, 100, 120, and 140 ms) and averaged across delays between the phase estimations of the envelopes and MEG data. A single MI value was generated per condition per ROI by concatenating all trials before calculating the MI (MEG and speech). Statistical analysis was performed per ROI per frequency band.

### Power analysis

Power analysis was performed to compare the MI results with absolute power changes, as any MI differences could be a consequence of signal-to-noise differences in the original data (which would be reflected in power effects). We first extracted the time-frequency representation for all conditions and ROIs separately. To do so, we performed a wavelet analysis with a width of 4, with a frequency of interest between 1 and 30 (step size of 1) and time of interest between −0.2 and 3.7 sec (step size of 0.05 sec). We extracted the logarithm of the power and baseline corrected the data in the frequency domain using a −0.3 and −0.1 sec window. For four different frequency bands (delta: 0.5-3.0 Hz; theta: 3.0-8.0 Hz; alpha: 8.0-15.0 Hz; beta: 15.0-25.0 Hz) we extracted the mean power in the 0.5-3.7 sec time window per task, condition and ROI. Again, our main analysis focusses on the delta band as that is where the main previous results were found [12], but we also report on the other bands for completeness. For each ROI we performed the statistical analysis on power as described in the behavioural analysis.

### Connectivity analysis

For the coherence analysis we repeated all pre-processing as in the power analysis, but separately for the left and right hemisphere (as we did not expect connections for PCA across hemispheres), after which we averaged the connectivity measure (using the Fourier spectrum and not the power spectrum). We used the debiased weighted phase lag index (WPLI) for our connectivity measure, which ensures that no zero-lag phase differences are included in the estimation (avoiding effects due to volume conduction). All connections between the three ROIs were investigated for the mean WPLI for the four different frequency bands in the 0.5-3.7 sec time window. Also in this case, the same statistical analysis was applied.

### Power control analysis

The reliability of phase estimations is influenced by the signal-to-noise ratio of the signal [25]. As a consequence, trials with generally high power have more reliable phase estimations compared to low power trials. This could influence any measure relying on this phase estimation, such as MI and connectivity [26, 27]. It is therefore possible that power differences between conditions lead to differences between connectivity or MI. To ensure that our reported effects are not due to signal-to-noise effects, we controlled any significant power difference between conditions for the connectivity and MI analysis. To do this, we iteratively removed the highest and lowest power trials between the mean highest and mean lowest of the two relevant conditions (either collapsing trials across tasks/conditions or using individual conditions). We repeated this until the original condition with the highest power had lower power than the other condition. Then we repeated the analysis and statistics, investigating if the effect of interest was still significant. The control analysis is reported along the main MI and connectivity sections.

## Results

### Behaviour

Overall task performance was above chance and participants complied with task instructions (Figure 1). We found a significant interaction between condition and task (F(2, 72.0) = 11.51, p < 0.001) as well as a main effect of task (F(2, 19.7) = 44.19, p < 0.001) and condition (F(2, 72.0) = 29.0, p < 0.001). We found that only for the word-combination (phrasal-level) task, the sentence condition had a significantly higher accuracy than the word list condition (t(54.0) = 6.97, p < 0.001). For the other two tasks, no significant condition effect was found (syllable: t(54.0) = 0.62, p = 1.000; word list: t(54.0) = 1.74, p = 0.176). Investigating the main effect of task indicated a difference between all tasks (phrase-syllable: t(18.0) = 3.71, p = 0.003; phrase-word: t(22.4) = −6.34, p < 0.001; syllable-word: t(19.2)=-8.67, p < 0.001).

**Figure 1.**
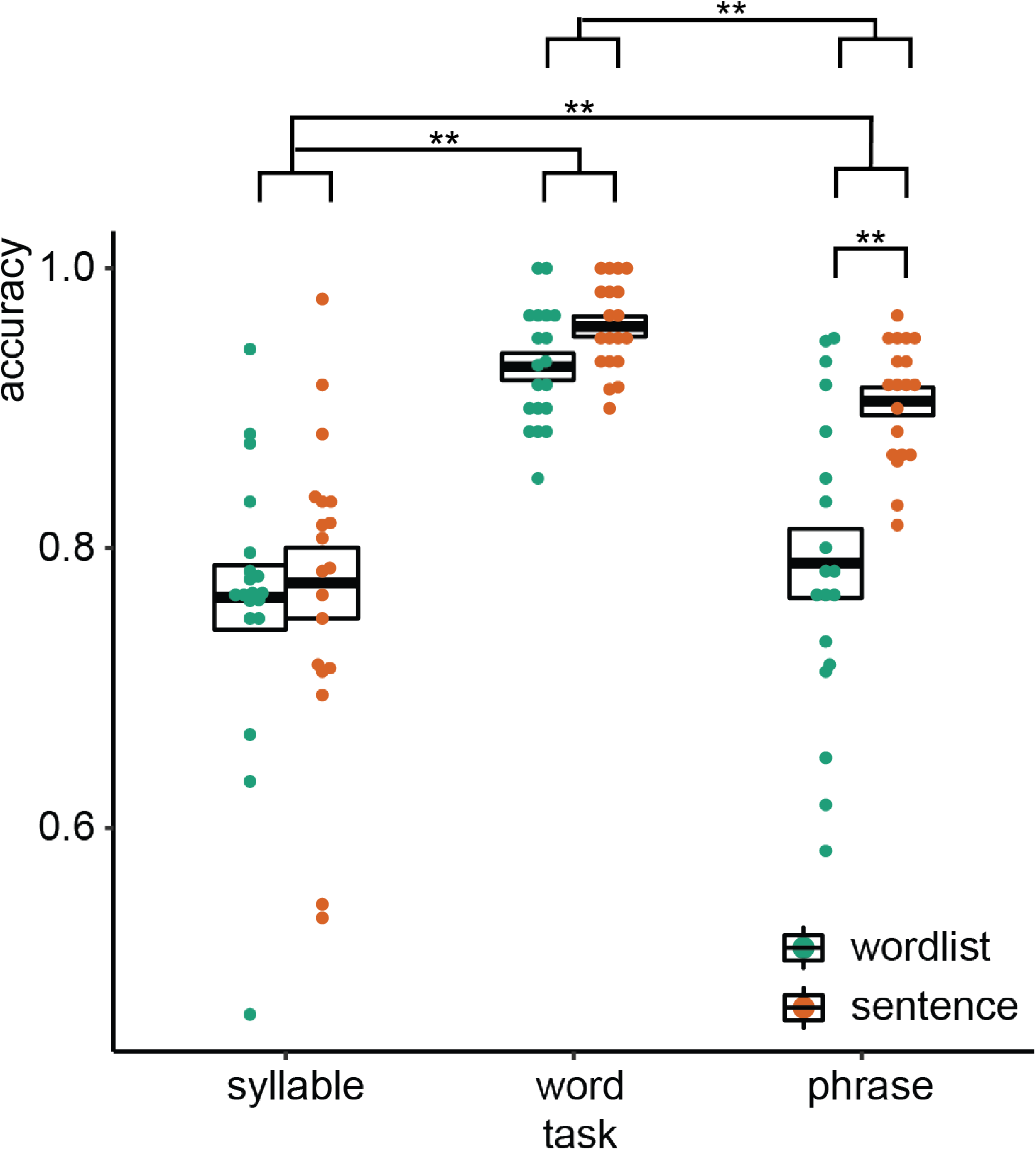
Behavioral results. Accuracy for the three different tasks. Double asterisks indicate significance at the 0.01 level.

### Mutual information

The overall time-frequency response in the three different regions of interest using the top-20 PCA components was as expected, with an initial evoked response followed by a more sustained response to the ongoing speech (Figure 2). From these regions-of-interest, we extracted mutual information in three different frequency bands (phrasal, word, and syllable). Here, we focus on the phrasal band as this is the band that differentiates word lists from sentences and showed the strongest modulation for this contrast in our previous study [12]. Mutual Information results for all other bands are reported in the supplementary materials.

**Figure 2.**
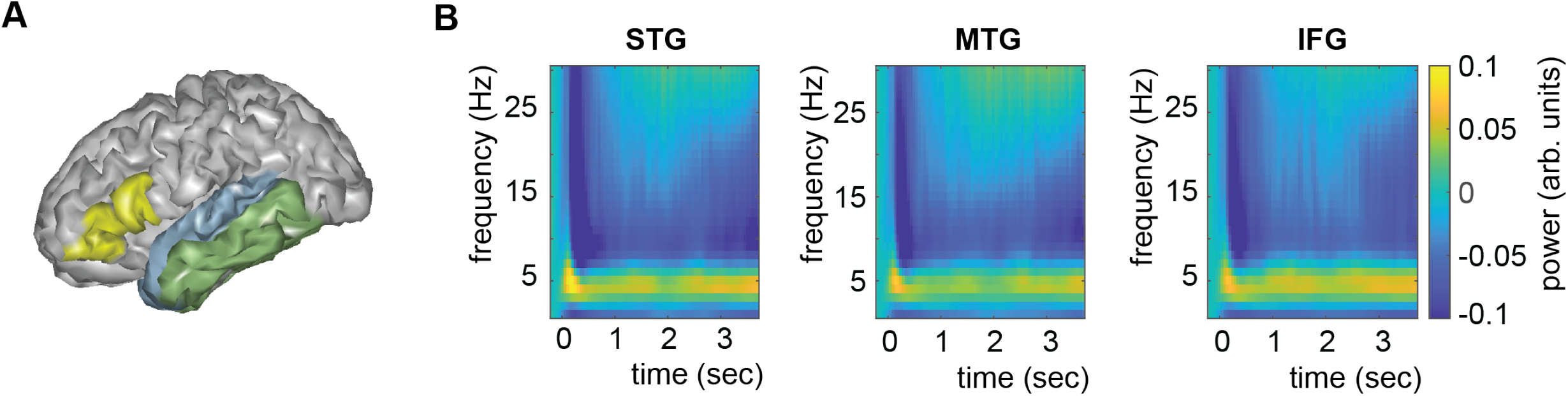
Anatomical regions of interests (ROIs). A) ROIs displayed one exemplar participant surface. B) Timefrequency response at each ROI. STG = superior temporal gyrus. MTG = medial temporal gyrus. IFG = inferior frontal gyrus.

For the phrasal timescale in STG, we found significantly higher MI in the sentence compared to the word list condition (F(3,126) = 67.39, p < 0.001; Figure 3). No other effects were significant (p > 0.1). This finding paralleled the effect found in Kaufeld et al., [12]. For the MTG, we saw a different picture: Besides the main effect of condition (F(3,126) = 50.24, p < 0.001), an interaction between task and condition was found (F(3,126) = 2.948, p = 0.035). We next investigated the effect of condition per task and found for all tasks except the passive task a significant effect of condition, with stronger MI for the sentence condition (passive: t(126) = 1.07, p = 0.865; syllable: t(126) = 4.06, p = 0.003; word: t(126) = 5.033, p < 0.001; phrase: t(126) = 4.015, p = 0.003). For the IFG, we found a main effect of condition (F(3,108) = 21.89, p < 0.001) as well as a main effect of task (F(3,108) = 2.74, p = 0.047). The interaction was not significant (F(3,108) = 1.49, p = 0.220). Comparing the phrasal task with the other tasks indicated higher MI for the phrasal compared to the word task (t(111) = 2.50, p = 0.028). We also found a trend for the comparison between the phrasal and the syllable task (t(111) = 2.17, p = 0.064), as well as the phrasal and the passive task (t(111) = 2.25, p = 0.052).

**Figure 3.**
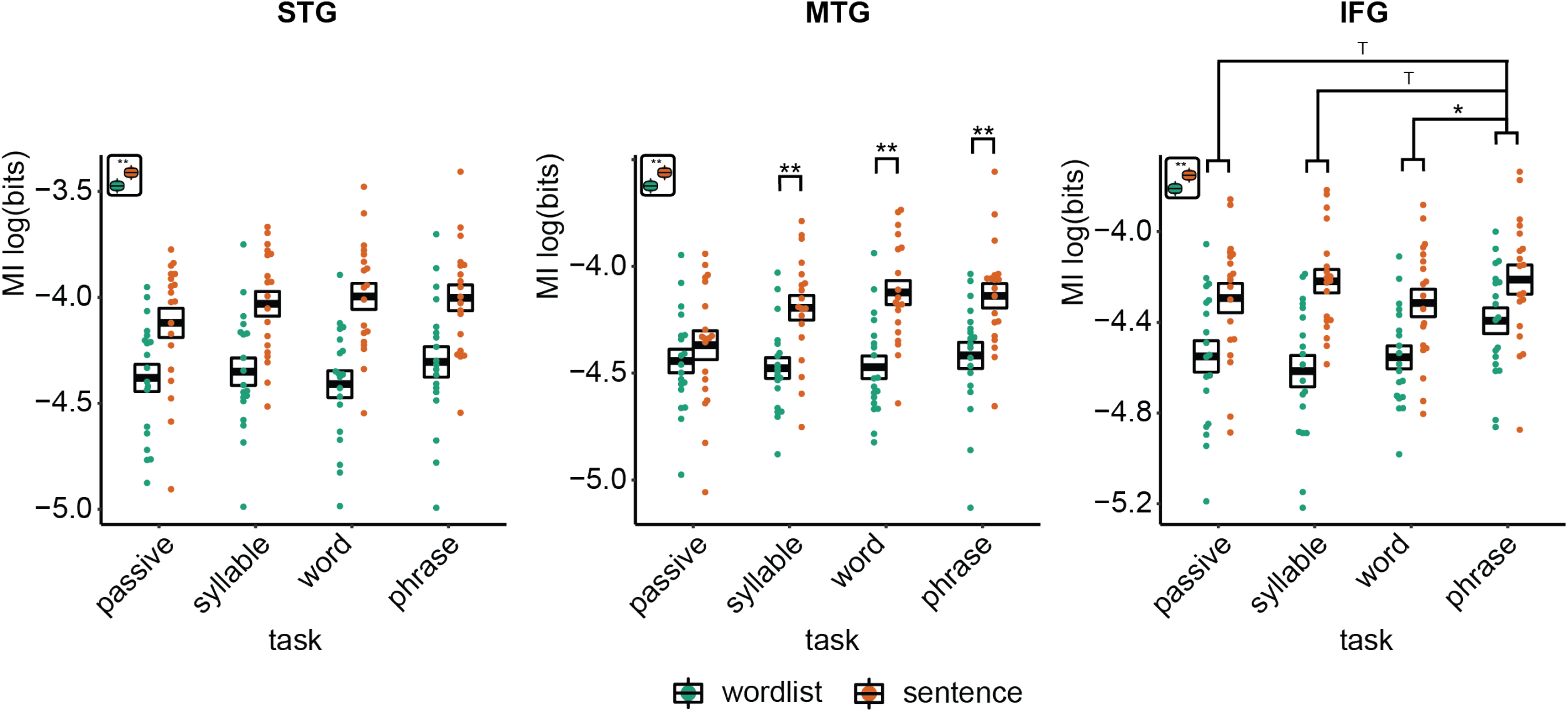
Mutual information (MI) analysis at the phrasal band (0.8-1.1 Hz) for the three different ROIs. Single and double asterisks indicate significance at the 0.05 and 0.01 level. T indicates trend level significance (p < 0.1). Inset at the top left of the graph indicate whether a main effect of condition was present (with higher MI for sentences versus wordlists).

For the word and syllable frequency bands no interactions were found (all p > 0.1; Supplementary Figure 1 and 2). For all six models there was a significant effect of condition, with stronger MI for word lists compared to sentences (all p < 0.001). The main effect of task was not significant in any of the models (p > 0.1; for the MTG syllable level there was a trend: F(3,126) = 2.40, p = 0.071).

When running the power control analysis, we did not find that significant effects in power differences (see next section; mostly due to main effects of condition) influenced our tracking results for any of the bands investigated.

### Power

We repeated the linear mixed modelling using power instead of MI to investigate if power changes paralleled the MI effects. For the delta band, we found for the STG a main effect of condition (F(1,18) = 6.11, p = 0.024) and task (F(3,108) = 3.069, p = 0.031). For the interaction we found a trend (F(3,108) = 2.620, p = 0.054). Overall sentences had stronger delta power than word lists. We found lower power for the phrase compared to the passive task (t(111) = 2.31, p = 0.045) and lower power for the phrase compared to the syllable task (t(111) = 2.43, p = 0.034). There was no significant difference between the phrase and word task (t(111) = 0.642, p = 1.00).

The MTG delta power effect overall paralleled the STG effects with a significant condition (F(1,124.94) = 12.339, p < 0.001) and task effect (F(3,124.94) = 4.326, p = 0.006). The interaction was trend significant (F(3,124.94) = 2.58, p = 0.056). Pairwise comparisons of the task effect showed significantly stronger power for the phrase compared to the passive task (t(128) = 2.98, p = 0.007) and lower power for the phrase compared to the syllable task (t(128) = 3.10, p = 0.024). The passive-word comparison was not significant (t(128) = 2.577, p = 0.109). Finally, for the IFG we only found a trend effect for condition (F(1,123.27) = 4.15, p = 0.057), with stronger delta power in the sentence condition.

The results for all other bands can be found in the supplementary materials (Supplementary figure 3–5). In summary, no interaction effects were found for any of the models (all p > 0.1). In all bands, power was generally higher for sentences than for word lists. Any task effect generally showed stronger power for the lower hierarchical level (e.g. generally higher power for passive versus phrasal tasks).

### Connectivity

Overall connectivity patterns showed the strongest connectivity in the delta and alpha frequency band (Figure 5). In the delta band, we found a main effect of task for the STG-IFG connectivity (F(3, 122.06) = 4.1078, p = 0.008). Follow-up analysis showed a significant difference between the phrasal and passive task (t(125) = 3.254, p = 0.003). The other comparisons with the phrasal task were not significant. The effect of task remained significant even when correcting for power differences between the passive and phrasal task (F(1, 53.02) = 12.39, p < 0.001; note the change in degrees of freedom as only the passive and phrasal task were included in this mixed model as any power correction is done on pairs). Initially, we also found main effects of condition for the delta and beta band for the MTG-IFG connectivity (stronger connectivity for the sentence compared to the word list condition), however after controlling for power, these effects did not remain significant.

**Figure 4.**
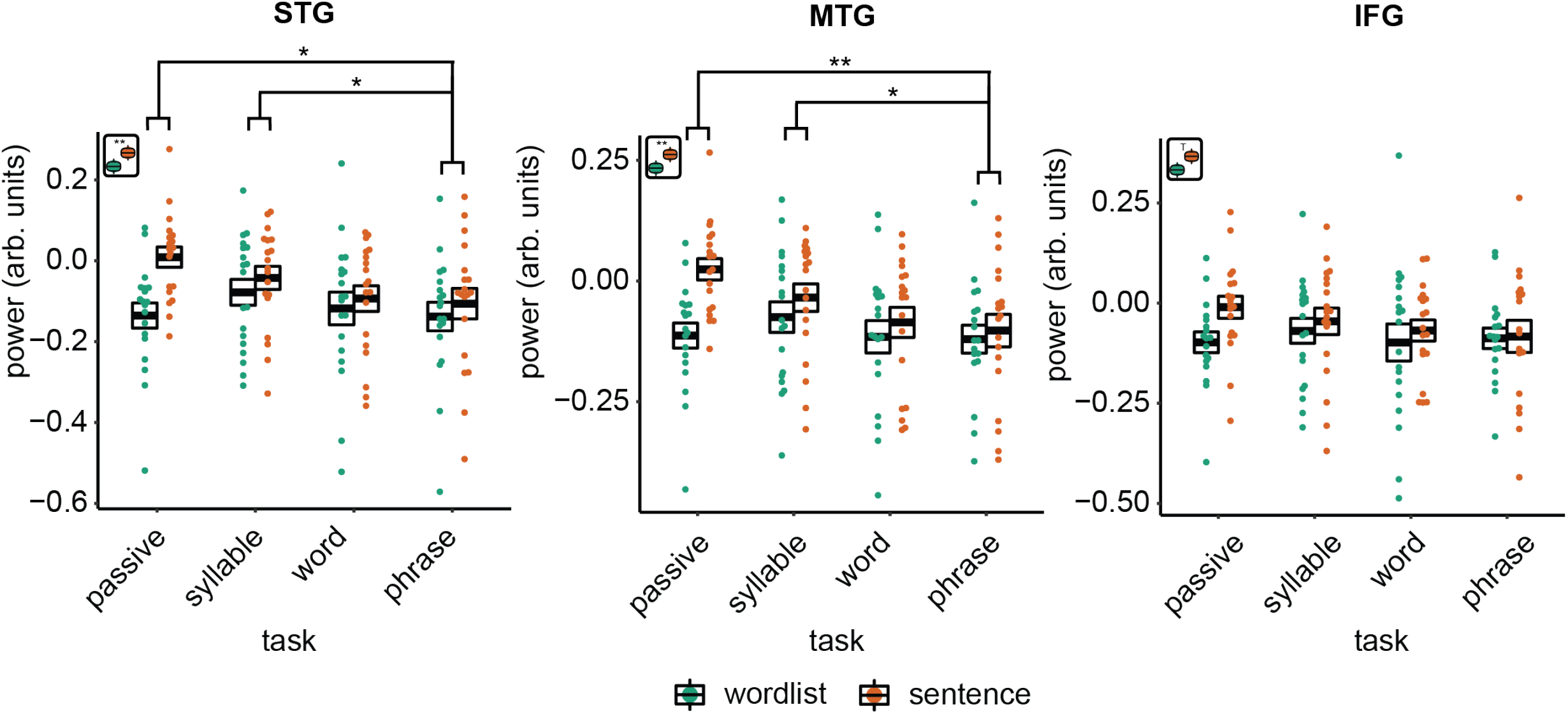
Power effects for the different ROIs. Single and double asterisks indicate significance at the 0.05 and 0.01 level. T indicates trend significance (p < 0.1) Inset at the left top of the graph indicate whether a main effect of condition was present (with higher activity for sentences versus wordlists).

**Figure 5.**
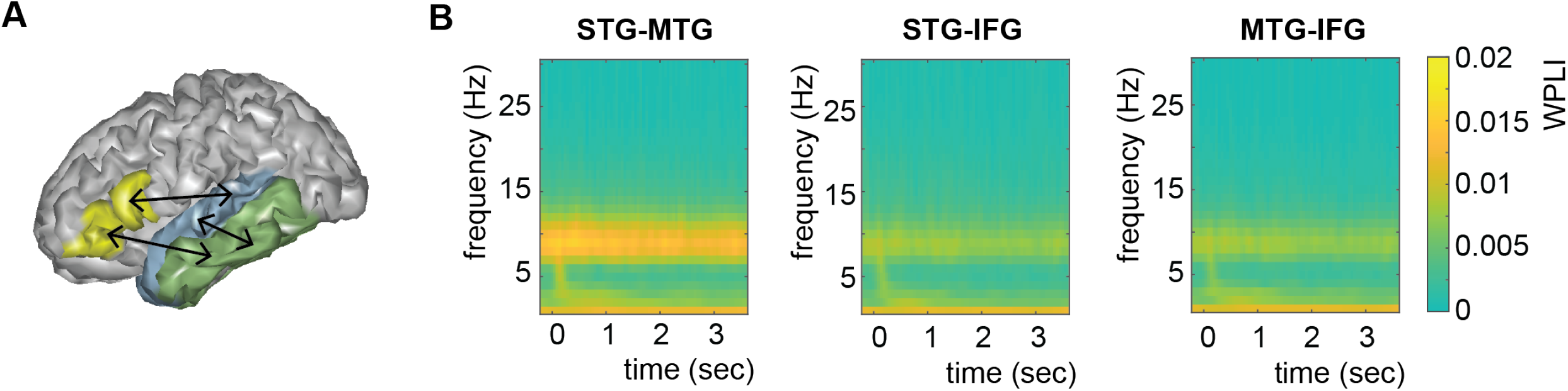
Connectivity pattern between anatomical regions of interests (ROIs). A) ROI connections displayed one exemplar participant surface. B) Time-frequency weighted phase-lagged index (WPLI) response at each ROI.

**Figure 6.**
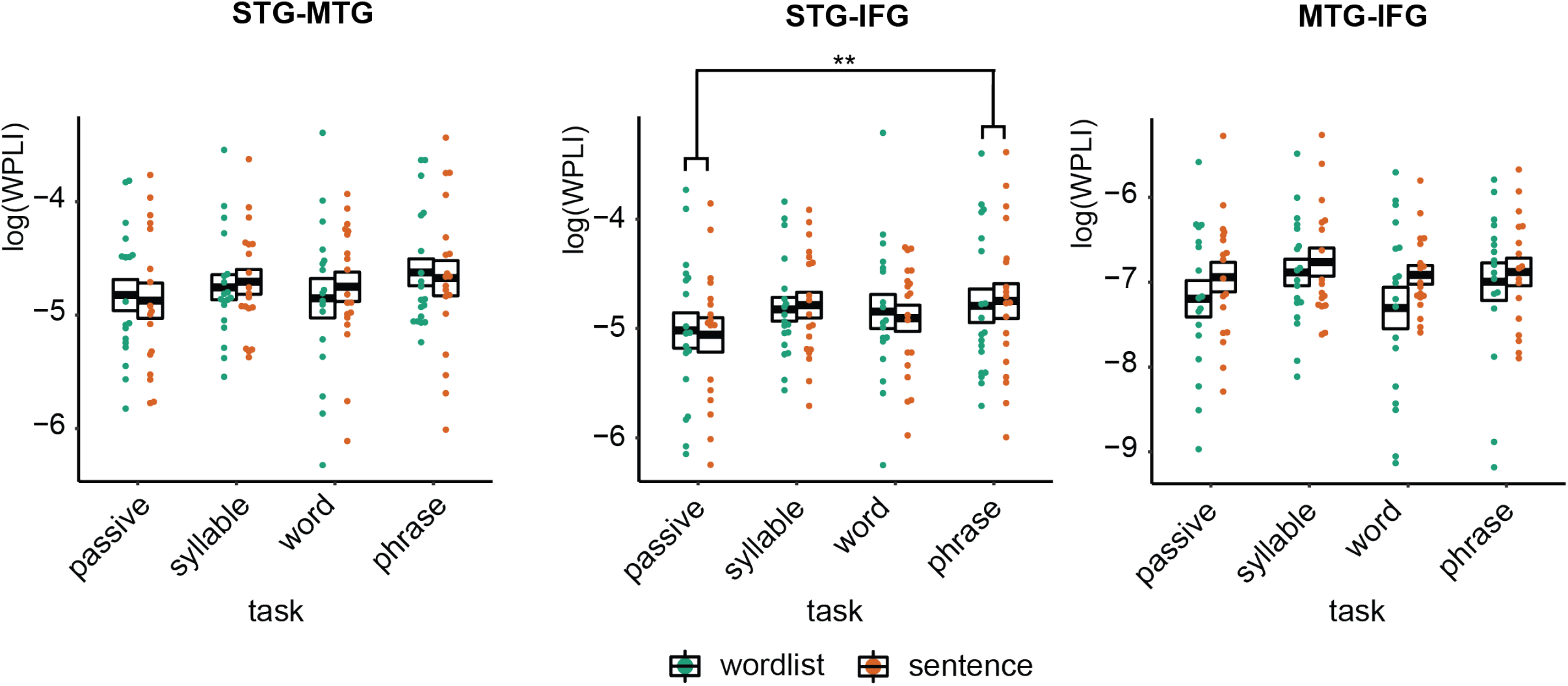
WPLI effects for the different ROIs. Single and double asterisks indicate significance at the 0.05 and 0.01 level after correcting for power differences between the two conditions (we plot the original data, not corrected for power, as we can only perform pairwise power and consequently data will be different for each control).

## Discussion

In the current study, we investigated the effects of ‘additional’ tasks on the neural tracking of sentences and word lists at temporal modulations that matched phrasal rates. Different nodes of the language network showed different tracking patterns. In STG, we found stronger tracking of phrase-timed dynamics in sentences compared to word lists, independent of task. However, in MTG we found this sentence-improved tracking only for active tasks. In IFG we also found an overall increase of tracking for sentences compared to word lists. Additionally, stronger phrasal tracking was found for the phrasal-level word-combination task compared to the other tasks (independent of stimulus type; note that for the syllable and passive comparison we found a trend), which was paralleled with increased IFG-STG connectivity in the delta band for the word combination task. This suggests that tracking at phrasal time-scales depends both on the linguistic information present in the signal, and on the specific task that is performed.

The findings reported in this study are in line with previous results, with overall stronger tracking of low frequency information in the sentences compared to the word list condition [12]. Crucially, for the stimuli used in our study it has been shown that the condition effects are not due to acoustic differences in the stimuli and also do not occur for reversed speech [12]. It is therefore most likely that our results reflect an automatic extraction of relevant phrase-level information in sentences, indicating the automatic processing of participants as they understand the meaning of the speech they hear using structural sentence information [2, 17, 28]. Overall, it did not seem that making participants pay attention to the temporal dynamics at the same hierarchical level through an additional task – instructing them to remember word combinations at the phrasal rate during word list presentation – could counter this main effect of condition.

Even though there was an overall main effect of condition, task did influence neural responses. Interestingly, the task effects differed for the three regions of interest. In the STG, we found no task effects, while in the MTG we found an interaction between task and condition. In the MTG increased phrasal-level tracking for sentences only occurred when participants were specifically instructed to perform an active task on the materials. It therefore seems that in MTG all levels of linguistic information are used to do an active language operation on the stimuli. This is in line with previous theoretical and empirical research suggesting a strong top-down modulatory response of speech processing in which predictions flow from the highest hierarchical levels (e.g. syntax) down to lower levels (e.g. phonemes) to aid language understanding [5, 29, 30]. As in the word list condition no linguistic information is present at the phrasal-rate, this information cannot be used to provide useful feedback for processing lower-level linguistic information. Instead, it could have been expected that the same type of increased tracking should have happened at the word-rate rather than the phrasal-rate for word lists (i.e., stronger word-rate tracking for word lists for the active tasks versus passive task). This effect was not found; this could either be attributed to different computational operations occurring at different hierarchical levels or to signal-to-noise/signal detection issues.

It is interesting that MTG, but not STG, showed an interaction effect. Both MTG and STG are strong hubs for language processing and have been involved in many studies which contrasted pseudo-words and words [31–33]. It is likely that STG does the lower-level processing of the two regions, as it is earlier in the cortical hierarchy, thereby being more involved in initial segmentation and initial phonetic abstraction rather than a lexical interface [31]. This could also explain why STG does not show task specific tracking effects; STG could be earlier in a workload bottleneck, receiving feedback independent of task, while MTG-feedback is recruited only when active linguistic operations are required. Alternatively, it is possible that either small differences in the acoustics are detected by STG (even though this effect was not previously found with the same stimuli [12]), or that our blocked designed put participants in a sentence or word list “mode” which could have influenced the state of these early hierarchical regions.

The IFG was the only region that showed an increase in phrasal-rate tracking specifically for the word-combination task. Note, however, that this was a weak effect, as the comparison between the phrase task and the syllable and passive task only reached a trend towards significance. Nonetheless, this effect is interesting for understanding the role of IFG in language. Traditionally, IFG has been viewed as a hub for articulatory processing [31], but its role during speech comprehension, specifically in syntactic processing, has also been acknowledged [1, 29, 34–36]. Integrating information across time and relative timing is essential for syntactic processing [2, 35, 37], and IFG feedback has been shown to occur in temporal dynamics at lower (delta) rates during sentence processing [38, 39]. However, it has also been shown that syntactic-independent verbal working memory chunking tasks recruit the IFG [35, 40–42]. This is in line with our findings that show that IFG is involved when we need to integrate across temporal domains either in a language-specific domain (sentences versus word lists) or for language-unspecific tasks (word combination versus other tasks). We also show increased delta-connectivity with STG for the only temporal-integration tasks in our study (i.e., the word combination task), independent of the linguistic features in the signal. Our results therefore support a role of the IFG as a combinatorial hub integrating information across time [43–45].

In the current study we investigated power as a neural readout during language comprehension from speech. This was both to ensure that any tracking effects we found were not due to overall signal-to-noise (SNR) differences, as well as to investigate task-and-condition dependent computations. SNR is better for conditions with higher power, which therefore leads to more reliable phase estimations, critical for computing MI as well as connectivity [25]. We will therefore discuss the power differences as well as their consequences for the interpretation of the MI and connectivity results. Generally, it seemed that there was stronger power in the sentence compared to the word list condition in the delta band. However, the pattern was very different than the MI pattern. For the power, the word list-sentence difference was the biggest in the passive condition. In contrast, for the MI there was either no task difference (in STG) or even a stronger effect for the active tasks (MTG; note that the power interaction was trend significant STG and MTG). We therefore think it unlikely that our MI effects were purely driven by SNR differences, and our power control analysis is consistent with this interpretation. Instead, power seems to reflect a different computation than the tracking, where more complex tasks generally lead to lower power across almost all tested frequency bands. As most of our frequency bands are on the low side of the spectrum (up to beta), it is expected that more complex tasks reduce the low-frequency power [46, 47]. It is interesting to observe that this did not reduce the connectivity for the delta band between IFG and STG, but rather increased it. It has been suggested that low power can potentially increase the available computational space, as it increases the entropy in the signal [48, 49]. Finally, in the power comparisons for the theta, alpha, and beta band we found stronger power for the sentence compared to the word list condition, which could reflect that listening to a natural sentence is generally less effortful than listening to a word list.

In the current manuscript we describe tracking of ongoing temporal dynamics. However, the neural origin of this tracking is unknown. While we can be sure that modulations in the phrasal-rate follow changes in the phrasal-rate of the acoustic input, it is unclear what the mechanism behind this modulation is. It is possible that there is stronger alignment of neural oscillations with the acoustic input at the phrasal rate [50, 51]. However, it could as well be that there is a phrasal time-scale or slower operation happening while processing the incoming input (which de facto is at the same time-scale as the phrasal structure occurring in the input). This operation, in response to stimulus input, could just as well induce the patterns we observe [52, 53]. Finally, it is possible that there are specific responses as a consequence of the syntactic structure, task, or statistical regularities occurring as specific events at phrasal time-scales [51, 54, 55].

It is difficult to decide on the most natural task in an experimental setting, that best reflects how we use language in a natural setting. This is probably why such a vast number of different tasks have been used in the literature. Our study (and many before us) indicates that during passive listening, we naturally attend to all levels of linguistic hierarchy. This is consistent with the widely accepted notion that the meaning of a natural sentence requires understanding the compositionality of words in a grammatical structure. For most research questions in language, it therefore is understandable to use a task that mimics this automatic natural understanding of a sentence. Here, we show that automatic understanding of linguistic information, and all the processing that this entails, cannot be countered to substantially change the consequences for neural readout, even when explicitly instructing participants to pay attention to particular time-scales.

## Supplementary figures

**Supplementary Figure 1.**
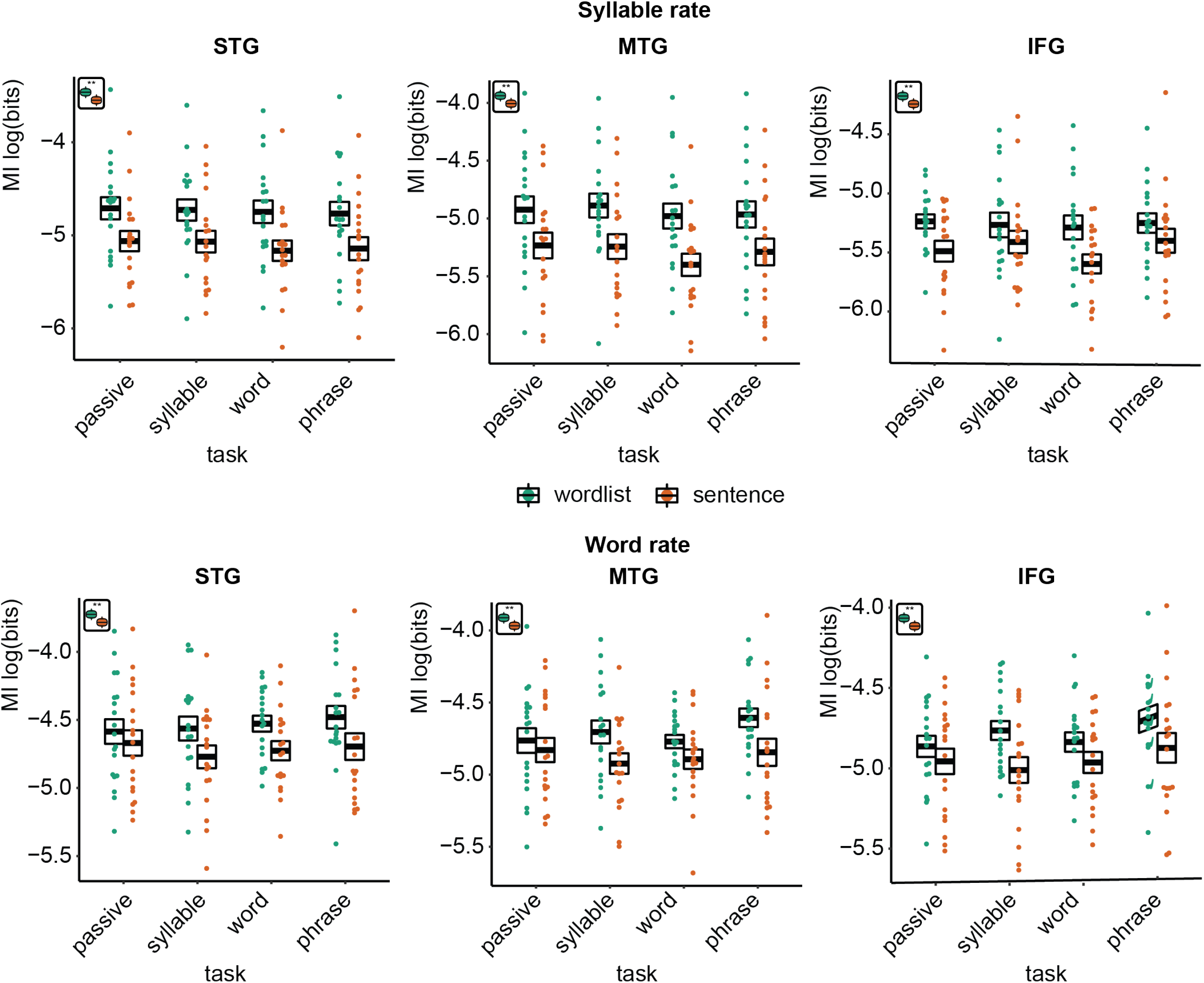
Mutual information (MI) analysis at the syllable (3.5-5.0 Hz) and word rate (1.9-2.8 Hz) for the three different ROIs. Double asterisks indicate significance at the 0.01 level. Inset at the top left of the graph indicate whether a main effect of condition was present (with higher MI for wordlists versus sentences).

**Supplementary figure 2.**
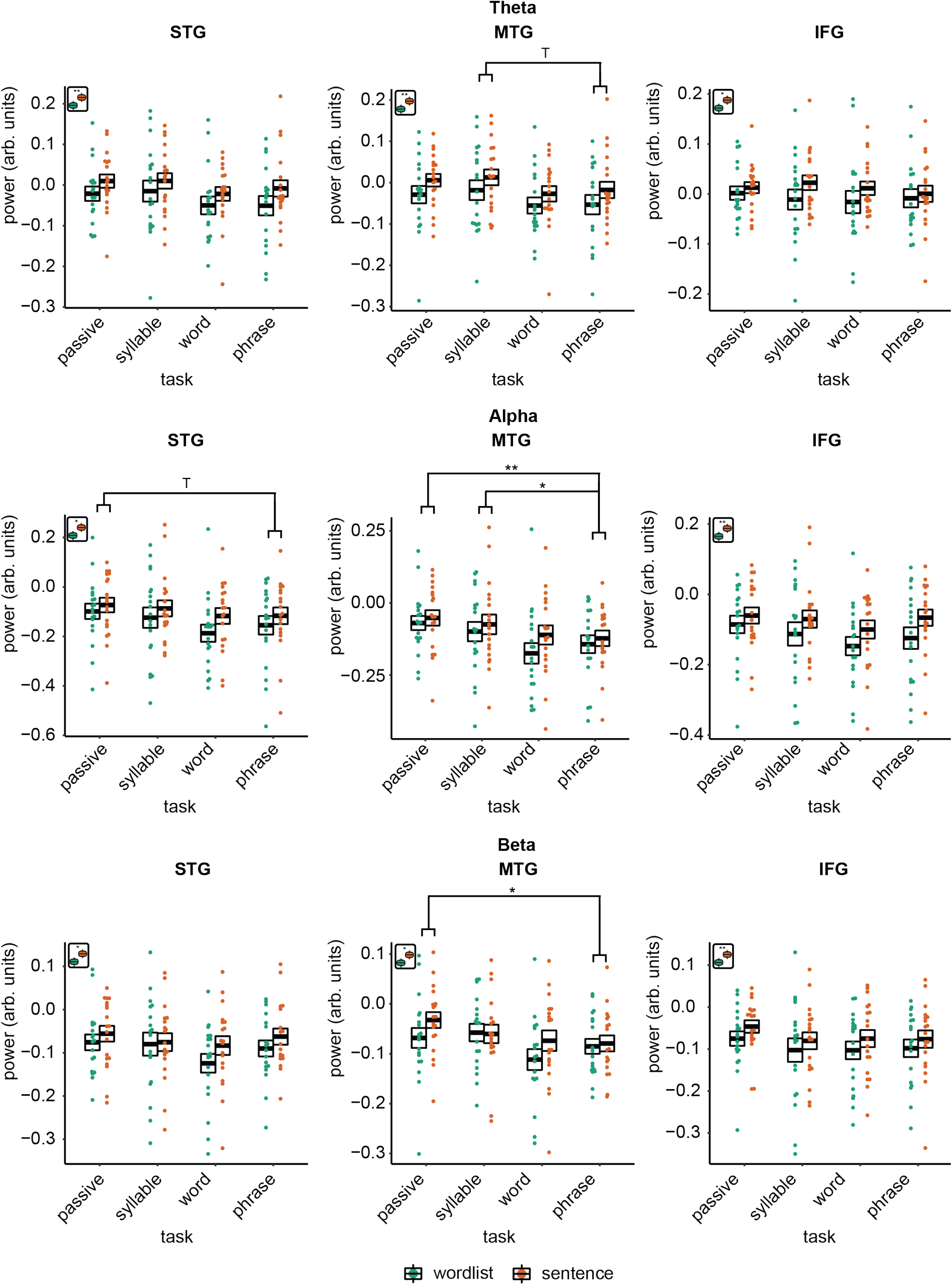
Power effects for the different ROIs and different bands. Single and double asterisks indicate significance at the 0.05 and 0.01 level. T indicates trend significance (p < 0.1) Inset at the top left of the graph indicate whether a main effect of condition was present (with higher activity for sentences versus wordlists).

**Supplementary figure 3.**
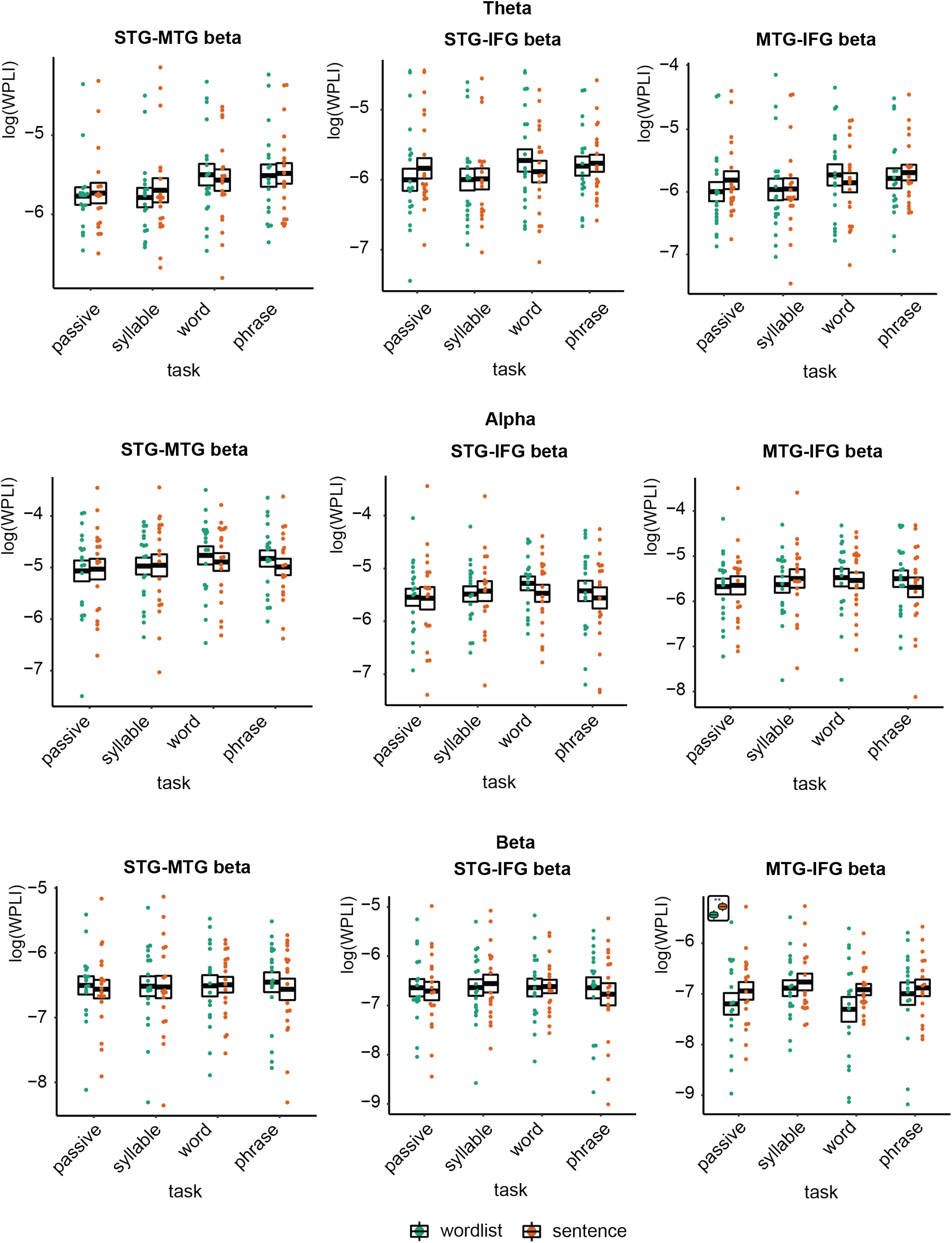
WPLI effects for the different ROIs and different bands. Connectivity is displayed before correcting for power differences. None of the effects survived correcting for power differences.

## Acknowledgments

AEM was supported by the Netherlands Organization for Scientific Research (NWO; grant 016.Vidi.188.029), and a Max Planck Research Group and a Lise Meitner Research Group “Language and Computation in Neural Systems” from the Max Planck Society.

## Notes

### Competing Interest Statement

The authors have declared no competing interest.

